# Forward planning driven by context-dependent conflict processing in anterior cingulate cortex

**DOI:** 10.1101/2021.07.19.452905

**Authors:** Florian Ott, Eric Legler, Stefan J. Kiebel

## Abstract

Forward planning is often essential to achieve goals over extended time periods. However, forward planning is typically computationally costly for the brain and should only be employed when necessary. The explicit calculation of how necessary forward planning will be, is in itself computationally costly. We therefore assumed that the brain generates a mapping from a particular situation to a proxy of planning value to make fast decisions about whether to use forward planning, or not. Moreover, since the state space of real world decision problems can be large, we hypothesized that such a mapping will rely on mechanisms that generalize sets of situations based on shared demand for planning. We tested this hypothesis in an fMRI study using a novel complex sequential task. Our results indicate that participants abstracted from the set of task features to more generalized control contexts that govern the balancing between forward planning and a simple response strategy. Strikingly, we found that correlations of conflict with response time and with activity in the dorsal anterior cingulate cortex were dependent on context. This context-dependency might reflect that the cognitive control system draws on category-based cognition, harnessing regularities in control demand across task space to generate control contexts that help reduce the complexity of control allocation decisions.

## Introduction

Many decisions have far-reaching consequences for the future, as they affect both internal bodily and external environmental states, in turn often conditioning potential future actions. Therefore, to achieve any long-term goals, people have to consider the future in some way. This can be achieved by planning multiple steps into the future to estimate the effects of potential action sequences (K. J. Miller et al., 2020; Simon et al., 2011; Tolman, 1948). However forward planning comes at a cost of using time and cognitive capacities, therefore people should only plan ahead when the benefits outweigh the costs and rely on fast and frugal strategies otherwise (Gershman et al., 2015; Gigerenzer et al., 2011; Kool et al., 2017; Lieder et al., 2020; Shenhav et al., 2013; Shenhav et al., 2017).

An intriguing question is how the brain controls when to engage in forward planning and when to use simpler strategies. Planning is often seen as one of the core functions of cognitive control (M. M. Botvinick et al., 2014; Goschke, 2013; E. K. Miller et al., 2001) and therefore the neural mechanisms involved in the regulation of cognitive control might be similarly involved in the regulation of planning. A classic hypothesis proposes that the dorsal anterior cingulate cortex (dACC) plays a central role in cognitive control by monitoring processing conflicts that serve as a signal for the need for additional control (M. M. Botvinick et al., 2001). Empirical evidence supports the involvement of the dACC in conflict processing in response interference tasks (Kerns et al., 2004; E. H. Smith et al., 2019), value-based decision making (Pochon et al., 2008) and recently in tasks that require people to plan multiple steps into the future (Economides et al., 2015; Korn et al., 2018; Schwartenbeck et al., 2015).

In multi-step tasks, however, an intricate computational problem becomes apparent. How can the brain control the use of forward planning in a way that maximizes long-term benefits, without having to compute these benefits by forward planning beforehand? One solution to this paradox might be for people to generate a mapping from a particular situation to a proxy of the value of planning that allows them to quickly access the planning values later (D. G. Lee et al., 2021; Lieder et al., 2018). Moreover, because the state space for real-world decision problems can be very large, it is unlikely that people learn a value for every possible combination of states. Rather, they might use certain task features to generalize clusters of states into particular contexts for which values are learned (Lieder et al., 2018).

Here, we tested this principle in an fMRI study using a novel sequential decision making task. In the task participants had to plan ahead to earn points by accepting offers while managing a limited energy budget. Importantly, we designed the task such that situations with different levels of the demand for planning occurred. With 448 possible combinations of task features and four different offers participants could choose from, our task was quite complex. We therefore assumed that participants used a simplified representation of planning value during control allocation decisions. An initial analysis of choice frequencies showed that participants used a repetitive choice pattern for two of the options, while responses were more balanced for the two other offers. From these choice patterns, we hypothesized that participants generated two different groups of offer-dependent representations of planning value. We refer to these two groups as control contexts (or context for short), with one context coding for a high a priori need for planning and the other context coding for a low a priori need for planning. To further test the control context hypothesis, we analysed response times and fMRI data using a specific conflict measure as a proxy for the value of forward planning. We found that correlations of conflict with response time and with BOLD-activity in the dACC were dependent on the context. Our results provide initial evidence for a mechanism by which the brain harnesses regularities in the value of planning across tasks space to construct control contexts that facilitate efficient allocation of control in complex tasks. Future research should further develop and confirm these initial findings by testing formal models of arbitration which incorporate structured representations of planning value.

## Methods

### Participants

Forty participants took part in the experiment (22 women, mean age = 24.4, SD = 4.6). Reimbursement was a fixed amount of 14€ or class credit plus a performance-dependent bonus (mean bonus = 6.62€, SD = 0.39). The bonus was calculated as a linear function of the accumulated points in the experiment. The study was approved by the Institutional Review Board of the Technische Universität Dresden and conducted in accordance to ethical standards of the Declaration of Helsinki. All participants were informed about the purpose and the procedure of the study and gave written informed consent prior to the experiment. All participants had normal or corrected-to-normal vision.

### Data availability

Data and analysis code used in this article is publicly available at https://doi.org/10.5281/zenodo.5112965. The repository includes: raw behavioural data and fMRI statistical maps underlying Figures 4 and 5; source code for reproducing Figures 2, 3, S2, and S3; source code implementing the model fitting, validation and comparison procedures.

### Experimental task

To investigate the context dependence of people’s propensity to plan ahead, we designed a novel open-ended sequential decision task in which participants had to accumulate as many points as possible. We induced a necessity for planning by introducing a limited but replenishable energy resource, which was required to accept offers. Planning was further encouraged by introducing variation around the energy cost of accepting and by explicitly informing participants about how these costs would change in the future. Unlike in experiments with a fixed deadline (Economides et al., 2014; Ott et al., 2020; Schwartenbeck et al., 2015), the open-ended nature of the task provided for every trial an opportunity to plan ahead multiple steps into the future. Importantly, the task featured both situations in which planning was crucial to decide between the accept and reject option, as well as situations that could be sufficiently solved by a simple heuristic.

In detail, the temporal structure of the task comprised three levels, the single trial, the current segment, and the segment pair of the current and next segment. One segment consisted of four trials. In a single trial, participants could either accept or reject an offer (selected by either a left or right button press), where accepting the offer increased points by an indicated amount but decreases energy by one or two units, depending on condition, see below. Rejecting always increased energy by one. There were four equally probable offers, displayed as one, two, three or four trophies in the middle of the screen. Accepting an offer increased points by the respective number of trophies, thereby advancing the yellow point bar at the top of the screen (Fig 1A). The energy costs of accepting varied between one, in low-cost segments (LC), and two in high-cost segments (HC). The energy budget with a minimum of zero and a maximum of six was displayed as a blue bar at the bottom of the screen. Initial energy in the first trial of the experiment was three. If participants had maximum energy and chose to reject, no further energy was added, and the next trial started. If participants accepted an offer with too little energy, no points were awarded, a warning was displayed, and the next trial started. Participants were informed about the energy cost of the current and the future segment by two symbols in the bottom right corner. The left symbol informed about both the energy cost of the current segment and the current trial number in the segment. The right symbol informed about the energy cost of the future segment. One flash indicated a low-cost segment and two flashes a high-cost segment (Fig 1A).

**Figure 1.**
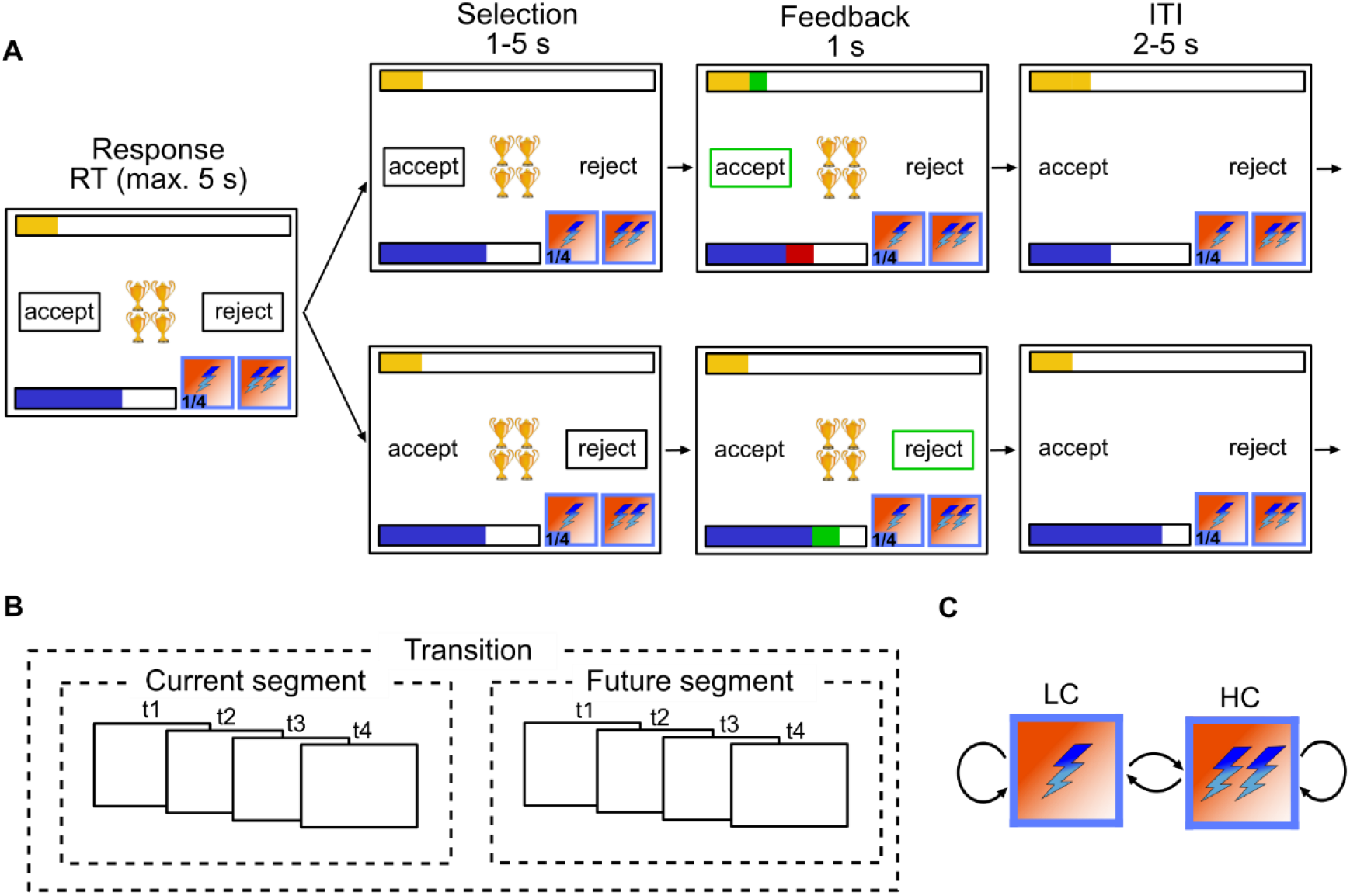
Experimental Task. **(A)** Timeline of a single trial. Participants could accept an offer (*O* = {1,2,3,4}) displayed in the middle of the screen to collect points (top yellow bar). Accepting an offer was associated with an energy cost. The current energy cost within a 4-trial-segment was indicated by the left symbol on the bottom right of the screen. One flash indicates an energy cost of 1 and two flashes an energy cost of 2. The right symbol on the bottom right indicates the energy cost of the next segment. Participants could choose the reject option to replenish the energy budget (bottom blue bar) by 1. **(B)** Temporal structure of the task: Four single trials formed a segment. Two consecutive segments formed a segment pair. **(C)** A segment could feature one of two different energy costs, low energy costs (LC) or high energy costs (HC). There were four different segment transitions LC/LC, LC/HC, HC/LC and HC/HC.

The experiment included a training session outside the MRI scanner (for task instructions see S1 Text) and three sessions inside. The training session comprised 144 trials across 36 segments with nine repetitions for each of the four possible transitions (Fig. 1C). The fMRI experiment comprised 240 trials across 60 segments with 15 repetitions per segment transition. On average, participants took 40 minutes to complete the fMRI experiment. The fMRI experiment was split into three sessions between which participants rested for two to three minutes without leaving the scanner. The sequence of segments and offers was pseudorandomized and identical for all participants. Segment sequences were generated such that each of the four segment transitions (Fig 1C) was sampled equally often. Similarly, offer sequences were generated such that the frequency of offers was balanced within segment transitions (raw behavioral data with all details about the offer sequences can be found at https://doi.org/10.5281/zenodo.5112965).

The timing of stimulus events in the fMRI experiment was as follows (see also Fig. 1A): each trial started with a fixation cross (0.5 seconds) in the middle of the screen to prepare participants for the upcoming decision. In the response phase, the offer appeared, and the choice options were surrounded by a frame to indicate that a decision is required. If participants did not respond within 5 seconds, they were timed out with a warning message, and the next trial began. In the selection phase (1-5 seconds, uniformly sampled and rounded to first decimal), the frame surrounding the unchosen option disappeared. In the feedback phase (1 second, fixed), energy or point changes were displayed, and the frame surrounding the chosen option turned green (or accordingly red if the energy budget was too low for accepting). In the intertrial interval (2-5 seconds, uniformly sampled and rounded to first decimal) choice options were unframed and the offer disappeared (Fig 1A). The training version of the stimulus was identical to the fMRI version except that there was no timeout for the response phase, there was not a selection phase and the intertrial interval was fixed to 1 second.

### Computational model of choice behaviour

We considered three different computational-cognitive models of how participants select their responses. Firstly, following one standard assumption about participants’ behaviour in such sequential decision making tasks, participants may have used forward planning across the current and the next segment (‘planning strategy’) to estimate the expected value of either accepting or rejecting (e.g. Kolling et al., 2014; Schwartenbeck et al., 2015). Secondly, in contrast, participants may have used some sort of heuristic to reduce the number of computations involved. An obvious choice for such a heuristic is the simple strategy to just base the accept/reject decision on the offer value (‘simple strategy’). For example, participants may simply always reject the lower offer values 1 and 2, and always accept offer values 3 and 4, provided there was enough energy left to accept the offer. Thirdly, we also considered a hybrid strategy of these two extremes, where participants may use a mixture between forward planning and simple strategy (‘hybrid strategy’). Note that our modelling approach relies on logistic regression and does not describe per se a process of how the brain may balance between forward planning and a simple strategy. Rather, the computational approach enables us to test for evidence that participants rely on (i) forward planning, (ii) a simple strategy or (iii) a mixture between these two extremes.

#### Planning strategy model (PM)

Clearly, if participants used a greedy strategy of accepting all offers in the first few trials of the task, they will quickly run out of energy and might not be able to accept better offers in future trials. Therefore, to maximize the accumulated points, one has to plan ahead, anticipating future actions, energy costs and reward opportunities. To implement such a planning strategy, we assumed a finite horizon until the end of the next future segment since participants were only explicitly informed about the energy costs of the current and the next future segment. As each segment had four trials each, this resulted in a horizon of maximally 8 trials and minimally 5 trials, i.e. when a participant has to select the decision for the fourth trial of the current segment. To derive a policy that maximizes expected reward over this horizon we formalised our task as a Markov Decision Process (MDP) (Puterman, 2014; Sutton et al., 2018). We define

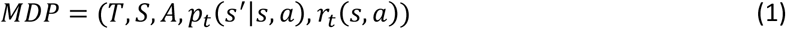

where *T* = {1,2,…,8} is the set of trials and *S* = *E* × *O* is the set of states with *E* = {0,1,…, 6} the set of possible energy levels and *O* = {1,2,3,4} the set of offer values. *A* = {*accept, reject*} is the set of actions.

Since participants successfully completed a training session and received detailed task instructions (S1 Text) prior to the main experiment, we assume that participants understood the rules of the task. In the model, this knowledge is represented by the transition probability *p_t_*(*s*′ = (*e′, o′*)|*s* = (*e, o*), *a*), which is the probability to transition to a new state *s*′ given the current state *s* (consisting of the offer value *o* and the current energy *e*) and the selected action *a*. Probabilities for allowed state transitions satisfy

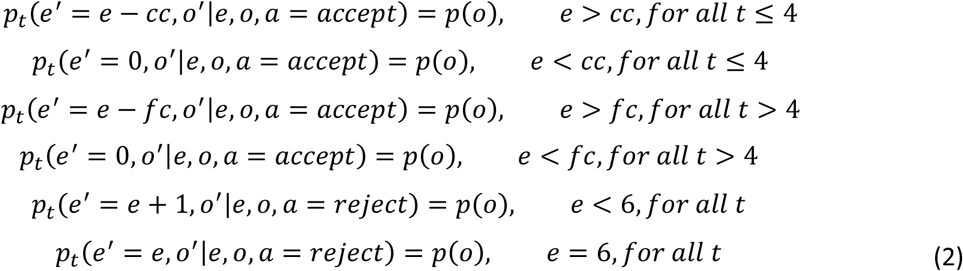

where *p*(*o*) is the discrete uniform distribution over the possible offer values, *cc* is the energy cost in the current segment (1 or 2) and *fc* is the energy cost in the future segment (1 or 2). We model the different segment transitions *LC* → *LC, HC* → *LC, LC* → *HC and HC* → *HC* as separate MDPs, substituting the respective values for *cc* and *fc*.

Immediate rewards, corresponding to the offer value, are generated upon successful acceptance. Formally, the reward function satisfies

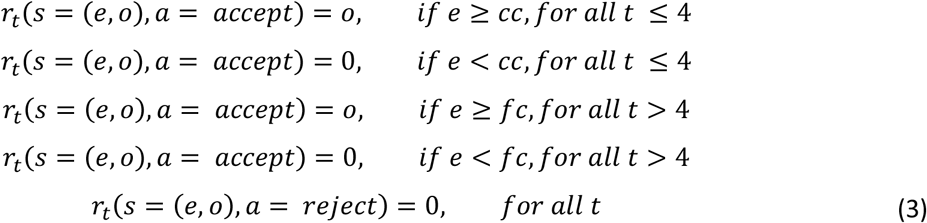

To determine the optimal policy that maximizes the expected reward over the current and future segment, the PM uses backward induction. The algorithm was implemented as follows:

1. Set *t* = 9 (if in the first trial of the segment) and define the state values after the decision in the final trial. To ensure that energy units left over after the eighth trial are considered to have utility, each remaining energy unit was multiplied by the quotient of the average offer value divided by the average energy costs.

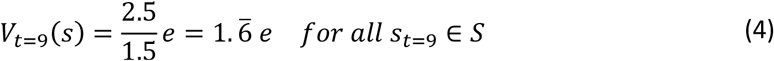
2. Set *t* = *t* – 1 and compute state-action values

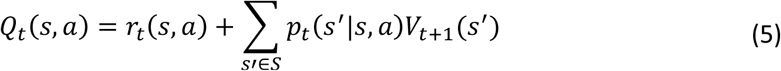
3. Update state values

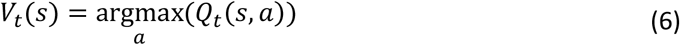
4. If t = 1 stop. Otherwise, continue with step 2

Action selection in the PM relies on a decision variable (DV) computed as the difference between the optimal state-action values

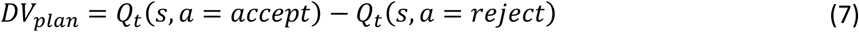

For a given state, positive values of DV indicate a greater long-term expected reward for accepting and negative values of DV indicate greater long-term expected reward for rejecting.

Using a logistic regression approach, we define the probability to accept as

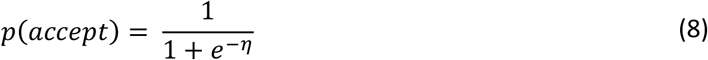

where

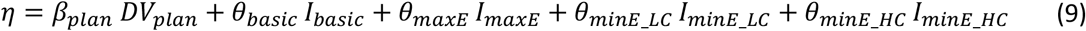

The planning weight *β_plan_* captures the influence of *DV_plan_* on choice behaviour. To allow for systematic deviations from behaviour prescribed by *DV_plan_*, we also included preference parameters *θ*. These preference parameters simply model a participant’s tendency to generally choose the accept (or the reject option). The parameter *θ_basic_* captures the preference in trials where participants had enough energy to accept and did not reach the maximum energy level (termed basic trials). We implemented this with a binary indicator variable *I_basic_* that equals one if the current trial was basic and zero if not. To model behaviour in trials with maximum or insufficient energy, we also included three bias parameters *θ_maxE_, θ_minE_HC_* and *θ_minE_LC_*. The first of these bias parameters *θ_maxE_* models the special case when participants had to choose on a trial with full energy. We expect this bias parameter to be generally positive because a further reject choice would not increase the energy further. For the other two bias parameters *θ_minE_HC_* and *θ_minE_LC_* we expect these to be generally negative, i.e. participants will reject an offer if they have insufficient energy. Subsets of these low- and max-energy trials are again selected by an appropriate binary indicator variable.

#### Simple strategy model (SM)

Since forward planning or other elaborate anticipatory schemes might incur considerable computational costs, participants may use a simple strategy, where action selection is only based on offer value. We define the decision variable for the SM as offer value centred across the four offer values 1 to 4:

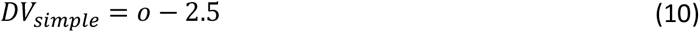

The probability to accept is defined in the same way as for the PM

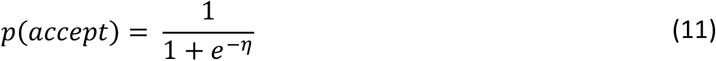

where

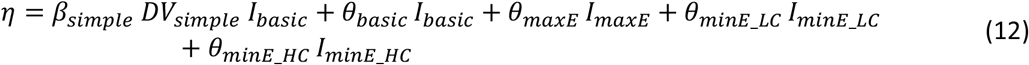

Here, the parameter *β_simple_* captures the influence of offer value on choice behaviour.

#### Hybrid strategy model (HM)

To cover the case that participants may choose based on both, expected long-term values and offer specific preferences, we use a hybrid strategy as a mixture of both planning and simple strategy. Such a hybrid strategy enables the decision maker to still use forward planning but mix this decision tendency with a simple strategy for each of the four offers. Note that we do not explicitly model arbitration and cannot identify which strategy dominates at any given time. However, the model enables us to test whether there is a mix of a simple and a planning strategy across trials. Like in the PM, *DV_plan_* is defined as the difference between the optimal state-action values

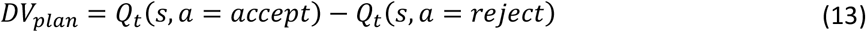

The probability to accept is defined as

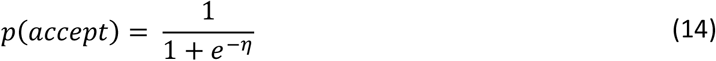

Where now

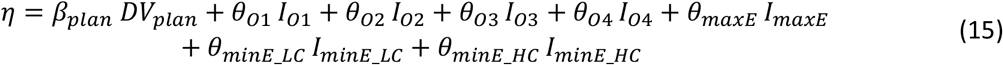

In addition to the planning weight *β_plan_* and the three bias parameters for extreme energy cases, the HM adds, as compared to the PM, four offer-specific preference parameters (*θ*_*O*1_, *θ*_*O*2_, *θ*_*O*3_, *θ*_*O*4_). The indicator variables (*I*_*O*1_, *I*_*O*2_, *I*_*O*3_, *I*_*O*4_) equal one if a specific offer was presented for basic trials (i.e. energy was neither at maximum nor too low to accept). In other words, in contrast to the PM, the four offer-specific bias parameters will indicate a relative dependence on the simple strategy. For example, a negative offer-specific parameter will indicate a participants’ preference to reject that specific offer.

### Model fitting and evaluation

#### Model fitting

Using a hierarchical Bayesian approach, we jointly estimated both participant- and group-level parameters. For the PM and the SM, *β* and *θ_basic_* were allowed to vary by participant. For the HM *β_plan_*, *θ*_*O*1_, *θ*_*O*2_, *θ*_*O*3_ and *θ*_*O*4_ were allowed to vary by participant. The parameters *θ_minE_LC_, θ_minE_HC_* and *θ_maxE_* were modelled as constant over participant. The participant parameters were drawn from a normal distribution with respective group parameters *μ* and *σ*. These group parameters were themselves modelled as draws from a weakly informative hyperprior distribution: *μ*~*Normal*(0,2) and *σ*~*HalfNormal*(0,2). A complete description of the models as Stan code can be found online (https://doi.org/10.5281/zenodo.5112965). We fitted models using Hamiltonian Markov Chain Monte Carlo as implemented in Stan (Carpenter et al., 2017) via the PyStan interface (Stan Development Team, 2018, Version 2.19.1.1). We obtained 4,000 samples from four chains of length 2,000 (1,000 warmup) from the posterior distribution over model parameters. The potential scale reduction factor on split chains 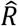 was calculated (Gelman et al., 1992), indicating convergence for all parameters 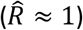.

#### Model comparison

We compared the predictive accuracy of the PM, SM and HM using leave-one-out cross-validation approximated by Pareto-smoothed importance sampling (PSIS-LOO) (Vehtari et al., 2017) as implemented in the python package ArViz (Kumar et al., 2019, Version 0.9.0). We obtained the expected log pointwise predictive density (elpd) and its standard error on the deviance scale (−2* elpd) and refer to this quantity as leave-one-out cross-validation information criterion (LOOIC). Lower values of LOOIC indicate better model fit.

#### Posterior predictions

To further assess whether the fitted models capture the observed behavioural pattern, we conducted posterior predictive checks using mixed predictive replication for hierarchical models (Gelman et al., 1996). To compute predictive replications we first sampled the group parameters (*μ* and *σ*) from the posterior and then sampled forty normally distributed participant-level parameters from these group parameters. Replicated accept-reject responses were generated for replicated participants and all trials by sampling from a Bernoulli distribution *response_rep ~ B*(*p*), where *p* is the response probability as defined in equations 8 and 9 (PM) or 12 (SM) or 15 (HM). The resulting array of replicated responses is of size *M* = *n_samples_* × *n_participnts_* × *n_trials_*. Individual dots in Figure 2A indicate choice proportions for 40 replicated participants averaged over 100 posterior samples. Bars represent averages over replicated participants and 100 samples, with error bars indicating between participant standard deviation. For the additional informal model validation in Figure 2C, we computed the proportion of replicated responses that matched the participant responses. Note that the set of stimulus configurations (energy, current segment type, future segment type, trial within segment) that participants visited during the task was identical to the stimulus configurations for which replicated responses were sampled under the three models (PM, SM, HM). A match was counted (match = 1, mismatch = 0) if the model’s response (accept or reject) was identical to a participant’s choice in a given trial.

**Figure 2.**
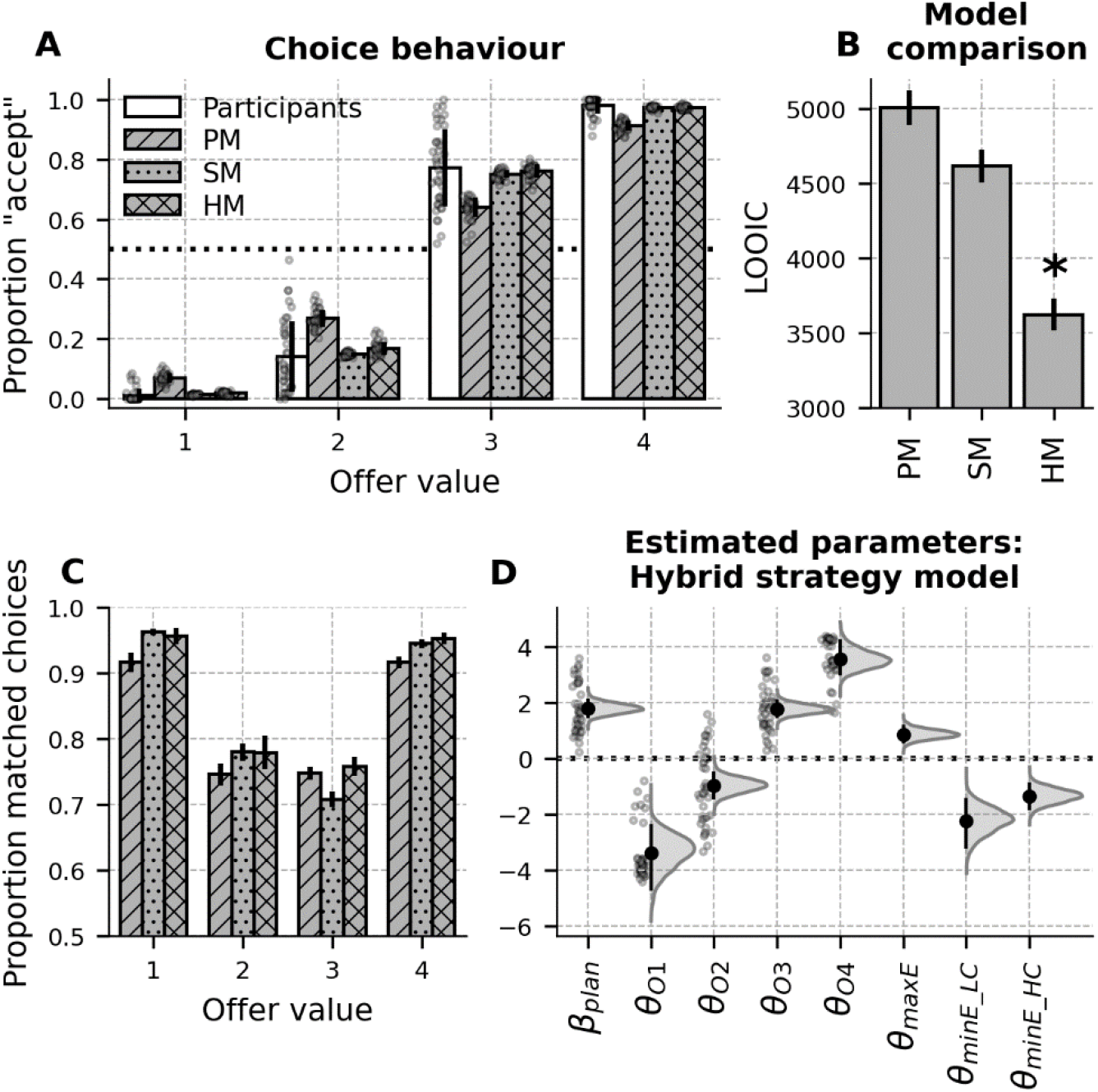
Choice behaviour and modelling results. **(A)** Plot of accept choices across offer values for participants and posterior predictions for the Planning strategy model (PM), simple strategy model (SM) and hybrid strategy model (HM). Bars represent the mean acceptance frequency across participants or across simulated participants. Trials in which energy was at max or too low to accept were excluded. Error bars represent standard deviation (SD). Dots represent individual participant data. The dotted line represents 50% chance level. **(B)** Model comparison of the PM, SM and HM. Error bars represent standard errors (SE) of the LOOIC. The asterisk indicates the winning model. **(C)** Proportion of simulated accept-reject choices that matched participant choices for the three models PM, SM and HM. Bars indicate averages over 200 posterior samples. The bar order and pattern is the same as in (A). Error bars represent SD. **(D)** Estimated parameters of the winning hybrid strategy model. Large black dots represent posterior means of group parameters with error bars depicting 95% credibility intervals. Grey curves represent kernel density estimates for the posterior distributions of group parameters. Semi-transparent small dots represent posterior means of participant-level parameter estimates.

#### Parameter recovery

To ensure that our model parameters are identifiable, we performed a parameter recovery analysis with the most complex model HM. We first generated simulated data using posterior means of the participant level parameters *β_plan_, θ*_*O*1_, *θ*_*O*2_, *θ*_*O*3_ and *θ*_*O*4_ and group level parameters *θ_minE_LC_, θ_minE_HC_* and *θ_maxE_*. Next, we refitted the model to the simulated data and compared the estimated parameters to the known data-generating parameters. We considered a known parameter recovered if its value was within the 95% posterior credibility interval (CI) of the re-estimated parameter. Results showed that both, participant level parameters (>99%) and all group level parameters, including those for *β_plan_*, *θ*_*O*1_, *θ*_*O*2_, *θ*_*O*3_ and *θ*_*O*4_ could be reliably recovered from the simulated data (for details see Jupyter Notebook at https://doi.org/10.5281/zenodo.5112965).

### Conflict and response time analysis

A key quantity for our analysis of response times (RT) and fMRI data was conflict.

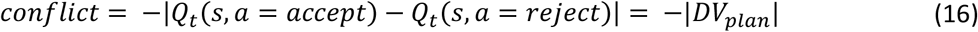

This corresponds to the similarity between long-term values for accepting and rejecting (see equation 7). If, for a given trial and task state, the action-value difference is small, conflict is large. Conversely, if the action-value difference is large, conflict is small. We consider conflict as a signal of choice difficulty, reflecting the need for elaborate information processing such as planning. We assume that participants do not calculate the conflict directly (which would require planning by itself), but that they have quick and frugal access to a proxy for the conflict (D. G. Lee et al., 2021).

We analysed response times using hierarchical Bayesian linear regression estimating group- and participant-level parameters simultaneously. We modelled log RT as the linear function

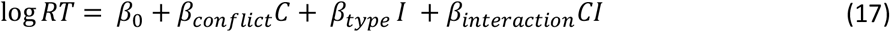

where *C* is conflict (Eq. 16), *I* is a binary indicator variable that equals one if the current offer was 2 or 3 (which we call in the following intermediate) and zero if the current offer is 1 or 4 (which we call in the following extreme). This classification into intermediate and extreme offers was based on participants’ choice behaviour (Fig. 2A). CI models the interaction between offer type and conflict. The participant-level intercept *β*_0_ and parameters *β_conflict_, β_type_* and *β_interaction_* were normally distributed with group parameters *μ* and *σ*. We gave these group parameters a weakly informative hyperprior: *μ~Normal*(0,10) and *σ~HalfNormal*(0,10). Models were fit in Stan via PyStan using Hamiltonian Markov Chain Monte Carlo. We obtained 2,000 posterior samples from four chains of length 1,000 (500 warmup). The potential scale reduction factor on split chains 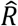 was calculated, indicating convergence for all parameters 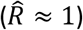. We generated linear predictions of log RT using 2,000 posterior samples of the group hyper parameters (*μ*_0_, *μ_conflict_, μ_type_, μ_interaction_*) and exponentiated back to the original RT scale for better interpretability. The regression lines in Figure 3C correspond to the median across samples and shaded areas to the 95% interval. All trials that were not timed out (RT > 5s) were included in the analysis.

**Figure 3.**
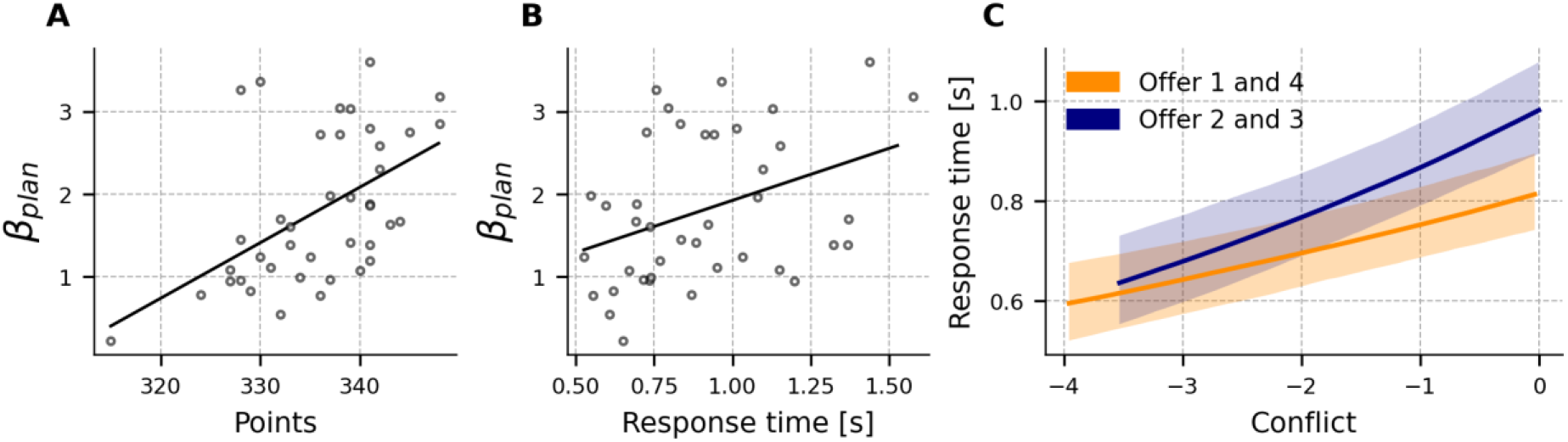
Correlation of fitted planning weight with behavioural markers and response time analysis. **(A)** Participant level posterior mean planning weight of the winning hybrid model versus accumulated points **(B)** Participant level posterior mean planning weight of the winning hybrid model versus mean response times. **(C)** Linear regression of log response times against choice conflict. Larger choice conflict was associated with an increase in response time, where the increase was significantly more pronounced during intermediate offers 2 and 3 compared to the extreme offers 1 and 4. Regression lines were computed from mean group hyperparameters. Shaded areas depict 95% credibility areas computed from 2,000 posterior samples of the group hyperparameters.

### fMRI acquisition and preprocessing

fMRI data were acquired on a 3 T MRI scanner (Siemens Magnetom Trio Tim, Siemens Medical Solutions, Erlangen, Germany) using a 32 channel head coil. On average, per participant, 942 volumes were acquired across three sessions, using a T2*-weighted echo-planar sequence (TR=2360 ms, TE=25 ms, flip angle=80°, FoV=192 mm). For each image, 48 axial slices of 2.5 mm were sampled in descending order. Field maps were acquired after each functional session (TR=532 ms, short TE=5.32 ms, long TE=7.78 ms). Structural data were acquired using a T1-weighted MPRAGE sequence (TR = 2400 ms, TE=2.19 ms, flip angle=8°, FoV=272 mm).

fMRI data were preprocessed and analysed using Statistical Parametric Mapping (SPM12) (Wellcome Trust Centre for Neuroimaging, London, UK). Functional images were unwarped using individual field maps generated by SPM’s field map toolbox (Hutton et al., 2002), slice time corrected, realigned to the first image of the session, spatially normalized to the MNI template using the unified segmentation approach (Ashburner et al., 2005) and smoothed with an 8mm full-width at half-max (FWHM) Gaussian kernel.

### fMRI Analysis

For each participant we specified and estimated a general linear model (GLM). Motivated by our behavioural results, we included one event regressor with response phase (see Fig. 1A) onsets for intermediate (2 and 3) offers and one event regressor with response phase onsets for extreme (1 and 4) offers. For each of these two event regressors, we included a parametrically modulated regressor with trial-wise conflict values. According to SPM’s default orthogonalisation setting, conflict values were mean-centred per condition. Additional regressors of no interest were included: an event regressor with response phase onsets for extreme energy trials (either maximum or insufficient energy) and an event regressor with response phase onsets of accept choices (to control for the effect of action). Onsets were modelled as stick function with duration 0. Regressors were convolved with the canonical HRF. We also included the 6 movement parameter vectors as nuisance regressors. Images of all three sessions were analysed together and a constant for each session was included in the design matrix. For each participant the following first level contrasts were computed: intermediate > extreme, extreme > intermediate, conflict_intermediate > conflict_extreme and a parametric effect of conflict averaged across intermediate and extreme trials. Finally, we performed one-sample t-tests on the contrast images of all participants to assess statistical significance on the group-level. Statistical parametric maps were initially thresholded with p = 0.001 (see Tables S2–S5). Voxels with a family-wise-error corrected p-value < 0.05 were considered significant.

dACC and dorsolateral prefrontal cortex (dlPFC) are commonly associated with conflict processing (E. K. Miller et al., 2001). Therefore, we defined a priori regions of interest (ROI), which we used for small volume correction (SVC). ROIs were defined using the WFU-Pickatlas software (Maldjian et al., 2003) with a dilation factor of 1. The ROI for the dACC encompassed dorsal Brodmann area (BA) 32, clipped at z = 18 in MNI-space (S1 Fig). The ROI for the dlPFC encompassed BA46 and BA9. Generated masks are available at https://doi.org/10.5281/zenodo.5112965.

## Results

### Behavioural results

#### Choice behaviour

We first identified situations in the task that could be classified as generally difficult or easy based on participants’ choice frequencies. An often repeated choice pattern indicates that a specific situation can be handled by simple response mechanisms, while a mixed response pattern of accept and reject indicates that more elaborate information processing may be required. Analysis of choice frequencies revealed an obvious pattern, showing that participants accepted offer 1 in only a few trials (mean = 1%, SD = 2%) and conversely accepted offer 4 in the majority of trials (mean = 98%, SD = 3%) (Fig. 2A). For offers 2 (mean = 14 %, SD = 12 %) and 3 (mean = 77 %, SD = 13 %), the choice behaviour was more balanced between accepting and rejecting. To further quantify the balance between accepting and rejecting, we computed the distance between choice frequencies and the 50% chance level and compared these distances across offer values. Distances from chance level were larger for offer 1 and 4 compared to offer 2 and 3 (pairwise Wilcoxon signed-rank tests, p < 0.001). There was no significant difference between offer 1 and offer 4 (Wilcoxon signed-rank test, p = 0.094). We also found that the distance from chance level was greater for offer 2 than for offer 3 (Wilcoxon signed-rank test, p < 0.001).

From this pattern, we hypothesised that participants might have treated the choice given an extreme offer 1 or 4 as generally easy and the choice given an intermediate offer 2 or 3 as generally hard. We hypothesized that this categorisation into what we call control contexts predetermines the actual planning investment in a given trial. In the following we will test this hypothesis and provide further insights using model-based analysis of choice behaviour, analysis of response times and analysis of fMRI BOLD-signals.

In addition to the offer value, we found, using logistic regression, that participants’ choice behaviour was also influenced by other task features (S1 Table). Participants chose more often the accept option if they had more energy units and likewise if the current energy cost of accepting was low (current segment = LC). Participants chose more often the accept option if the upcoming energy cost was high (future segment = HC), showing that participants considered information about the future segment when making a decision.

#### A combination of forward planning and simple offer specific preferences fitted behaviour best

We assumed that our task design motivated participants to use simple heuristics and forward planning in a situation-appropriate way. To test for the simultaneous presence of planning and simple heuristics we carried out a model-based analysis of choice behaviour. Three different strategies of how participants select their responses were considered. First, participants may have fully relied on forward planning across the current and the next segment to select actions that maximize the expected value (PM). Second, participants may have used a simple strategy just dependent on the offer value (SM). To illustrate the difference between these two strategies, let us consider a (for convenience deterministic) agent in the first trial of a segment, with three energy units, where the current and the future energy cost is 2 (segment pair HC/HC) and offer 3 is presented. A planning agent would reject the offer and replenish its energy reserves in order to be able to accept potential better offers in the future. In contrast, an agent following a simple strategy, e.g. who always accepts offers 3 and 4 and rejects offer 1 and 2, would accept the offer 3. We also considered a third alternative that participants use a mixture between planning and a simple strategy (HM) to achieve a good trade-off between the benefits and costs of the respective strategies depending on the current task situation.

We compared how well the three cognitive computational models fitted participants’ behaviour using leave-one-out cross validation. We found, as shown in Figure 2B, that the HM explained participant behaviour substantially better (LOOIC = 3622.5, SE = 108) than the PM (LOOIC = 5006.0, SE = 117.5) and the SM (LOOIC = 4613.2, SE = 109.4). This demonstrates that participants use both planning and simple heuristics throughout the task. To confirm this result, we also compared models on the participant level and found that the HM explained behaviour best for 33 (of N = 40) participants (S2 Fig). For 3 participants the SM and for 4 participants the PM explained behaviour best.

We also simulated posterior predictions for the three models and plotted acceptance frequencies across offer values (Fig 2A). Both, the HM and the SM, closely captured the behavioural pattern of participants, but acceptance frequencies of the PM were lower for offer 1 and 2 and higher for offer 2 and 3. These simulations are consistent with participants mixing forward planning with a simple reject-preference for offers 1 and 2 and an accept-preference for offers 3 and 4. As a further informal illustration of why the HM was superior to the SM, we computed the proportion of matches between participant choices and the simulated choices from the fitted models (Fig 2C). While SM shows high matching rates for offers 1, 2 and 4, the matching rate for offer 3 is decisively lower compared to the HM. Conversely, the PM has considerable lower matching rates than the HM for offers 1,2,4 but achieves a relatively high matching rate for offer 3. These results show that the SM particularly fails to account for participants choices for offer 3, presumably because participants engage in an increased amount of planning for offer 3 (see Fig. 2A, where the mean accept rate for offer 3 is closest to the 50% line among all four offers, i.e. offer 3 does not support a simple action selection strategy).

Parameter estimates of the HM demonstrate both evidence for forward planning and usage of a simple strategy as quantified by four offer-specific preferences (Fig 2D). We found a positive weight on the planned value difference (mean group parameter *β_plan_* = 1.79, 95% CI = [1.45, 2.15]). This indicates that participants, when making a decision, accounted for its future consequences. We also found preference parameters different from zero for all four offers. For offer one (mean group parameter *θ*_*O*1_ = −3.39, 95% CI = [−4.76, −2.37]) and offer two (mean group parameter *θ*_*O*2_ = −0.96, 95% CI = [−1.47, −0.47]), participants showed a preference for rejecting (indicated by negative parameter values). For offer three (mean group parameter *θ*_*O*3_ = 1.77, 95% CI = [1.43, 2.11]) and offer four (mean group parameter *θ*_*O*4_ = 3.56, 95% CI = [2.97, 4.28]), participants showed a preference for accepting (indicated by positive parameter values). We explicitly modelled special cases, where participants had either maximum energy (*θ_maxE_*) or not enough energy to accept (*θ_minE_LC_* and *θ_minnE_HC_*), see also Methods. As expected, participants showed a bias to accept in maximum energy trials and a bias to reject in low energy trials (see Fig 2D).

We further found evidence that participants who account for the long-term consequences of their actions by e.g. planning, earn more points. Post-hoc correlation analysis revealed that participants with a larger fitted planning parameter *β_plan_* of the winning hybrid model accumulated more points throughout the experiment (r = 0.521, p = 0.001, Fig 3A) and had slower average response times (r = 0.374, p = 0.017, Fig 3B).

#### Greater conflict-driven increase of response times in intermediate than extreme trials

Previous research suggested that the brain regulates the use of cognitive control based on the estimated value of control (Shenhav et al., 2013). Analogously, we assume that the brain uses similar value estimates when deciding about the degree of forward planning during sequential decisionmaking. Most importantly we hypothesized that, due to cost incurred by the computation of control values themselves, the brain uses a context-specific prior assumption about the general need for planning to minimize the “metacosts” of control decisions. To further test this hypothesis we analysed the relationship between response times as an indicator for the degree of planning and planning value (operationalised by a specific conflict measure, see methods). We expected that not only would larger conflicts generally lead to increased response times, but critically that this increase will be more pronounced for the intermediate offers 2 and 3, possibly reflecting context-specific planning activity driven by a context-specific evaluation of conflict.

Bayesian linear regression indeed showed that conflict was significantly more predictive for log RT for the intermediate offers 2 and 3 compared to the extreme offers 1 and 4 (group parameter *β_interaction_* = 0.04, 95% CI = [0.02 0.07]). We also found a significant positive main effect of conflict (group parameter *β_conflict_* = 0.08, 95% CI = [0.05 0.11]) and offer type (group parameter *β_is_intermediate_* = 0.18, 95% CI = [0.14 0.23]) on log RT. Figure 3C shows fitted regression lines on the untransformed RT scale. This shows that, the increase in computation time associated with conflict level is greater for intermediate offers than extreme offers.

### FMRI results

#### A frontal network is more activated in intermediate than extreme trials

A set of parietal and frontal regions, sometimes called the multiple demand network, are often activated during cognitively challenging tasks (Duncan, 2010; E. K. Miller et al., 2001). To relate to these well-established findings, we first tested to confirm where brain activity was higher during intermediate compared to extreme offers (Intermediate > Extreme) expecting to see greater activity in frontal areas related to planning and cognitive control. We indeed found significantly greater activity in dACC and right dlPFC (Fig 4A, Table 1, and S1 Table).

**Figure 4.**
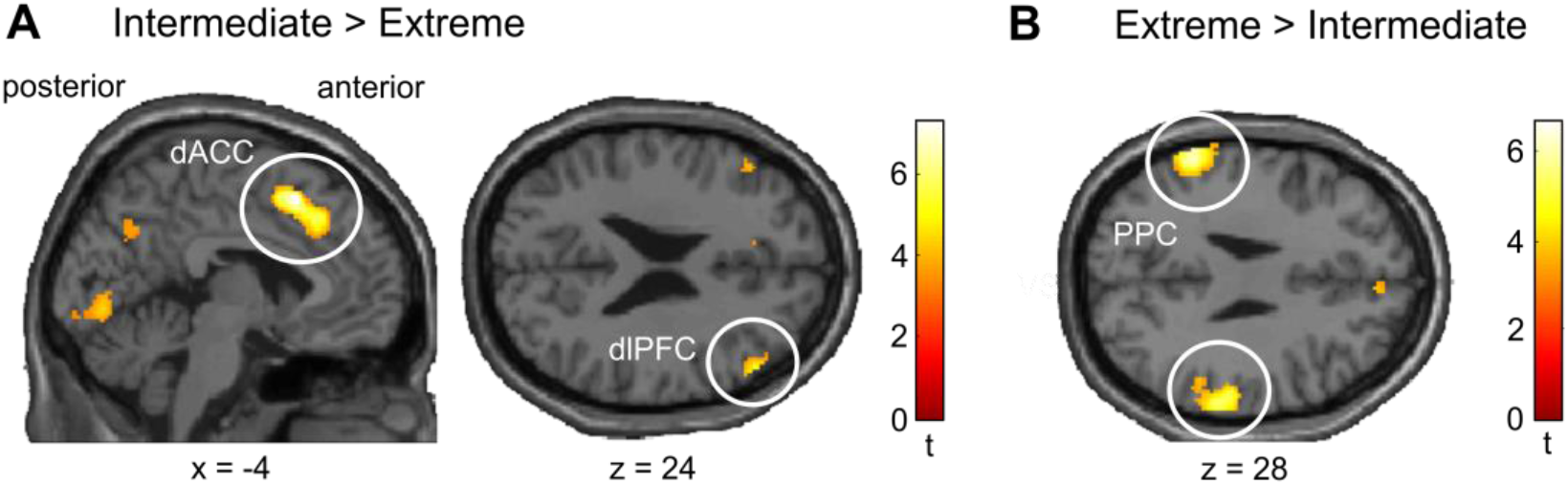
Different activation of dACC for intermediate versus extreme offers. **(A)** dACC and right dlPFC were significantly more activated during intermediate compared to extreme offers. **(B)** Bilateral PPC was significantly more activated during extreme compared to intermediate offers. Activations displayed at p < 0.001 uncorrected. See Table 1 for peak MNI-coordinates and statistics, significant at p<0.05 FWE corrected.

**Table 1.** Summary of fMRI Results.

|  | t | FWE p value |  | MNI coordinates |  |  |
| --- | --- | --- | --- | --- | --- | --- |
| Region |  | whole brain | SVC | x | y | z |
| Intermediate > Extreme |  |  |  |  |  |  |
| dACC | 7.26 | < 0.001 | < 0.001 | -4 | 16 | 52 |
| dIPFC R | 5.12 | 0.180 | 0.018 | 52 | 34 | 24 |
| Occipital L | 6.98 | < 0.001 |  | -18 | -92 | -10 |
| Occipital R | 6.38 | 0.007 |  | 20 | -84 | -8 |
| Extreme > Intermediate |  |  |  |  |  |  |
| PPC (BA 40) L | 6.68 | 0.003 |  | -60 | -44 | 38 |
| PPC (BA 40) R | 5.49 | 0.072 |  | 64 | -32 | 28 |
| Conflict |  |  |  |  |  |  |
| dACC | 4.82 | 0.290 | 0.007 | 10 | 24 | 44 |
| anterior Insula L | 5.62 | 0.041 |  | -30 | 24 | -2 |
| anterior Insula R | 6.50 | 0.004 |  | 38 | 20 | 0 |
| Conflict: Intermediate > Extreme |  |  |  |  |  |  |
| dACC | 4.32 | 0.744 | 0.030 | 8 | 2 | 40 |
| posterior MTG R | 5.69 | 0.040 |  | 56 | -64 | 6 |

Regions significant ( $p < 0.05$ FWE corrected) at the whole brain level or small volume correction (SVC) based on a priori regions of interest. Only clusters with more than 10 voxels were reported. BA, Brodmann area; L, left; R, right; dACC, dorsal anterior cingulate cortex; dlPFC, dorsolateral prefrontal cortex; Occipital, occipital lobe; VS, ventral Striatum; PPC, posterior parietal cortex; MTG, middle temporal gyrus.

We also tested where brain activity was greater during extreme versus intermediate trials (Extreme > Immediate) and found increased activity in bilateral posterior parietal cortex, where the cluster in the left hemisphere was significant at the whole brain corrected level (PPC; Fig 4B, Table 1 and S3 Table). Besides its role in sensory attenuation, posterior parietal cortex is also involved in sensorimotor transformations during decision making (Andersen et al., 2009). This suggests that participant decisions for extreme offers might be related to low-level sensorimotor processes, coupling simple stimulus cues to actions. A network including left ventral Striatum (VS), posterior cingulate cortex (PCC) and bilateral Amygdala emerged at a lower threshold (see S3 Table). These regions have been shown to encode value information during reward-based choice (Bartra et al., 2013). Higher activation in these areas during extreme trials might indicate an increased salience of offer value information instigating a simple response strategy based on offer-specific preferences. However, this idea requires further research.

#### Context-dependent conflict processing in dACC

As a confirmatory analysis of previous findings implicating the dACC in the monitoring of various signals to evaluate the need for additional control (e.g. conflict, Shenhav et al., 2013), we also tested for the effect of conflict averaged across conditions. In accord with this previous research, we found a significant positive correlation with BOLD-activity in the dACC (Fig 5A, Table 1 and S4 Table). We also found a significant positive effect of conflict in bilateral anterior Insula (Fig 5A, Table 1 and S4 Table).

**Figure 5.**
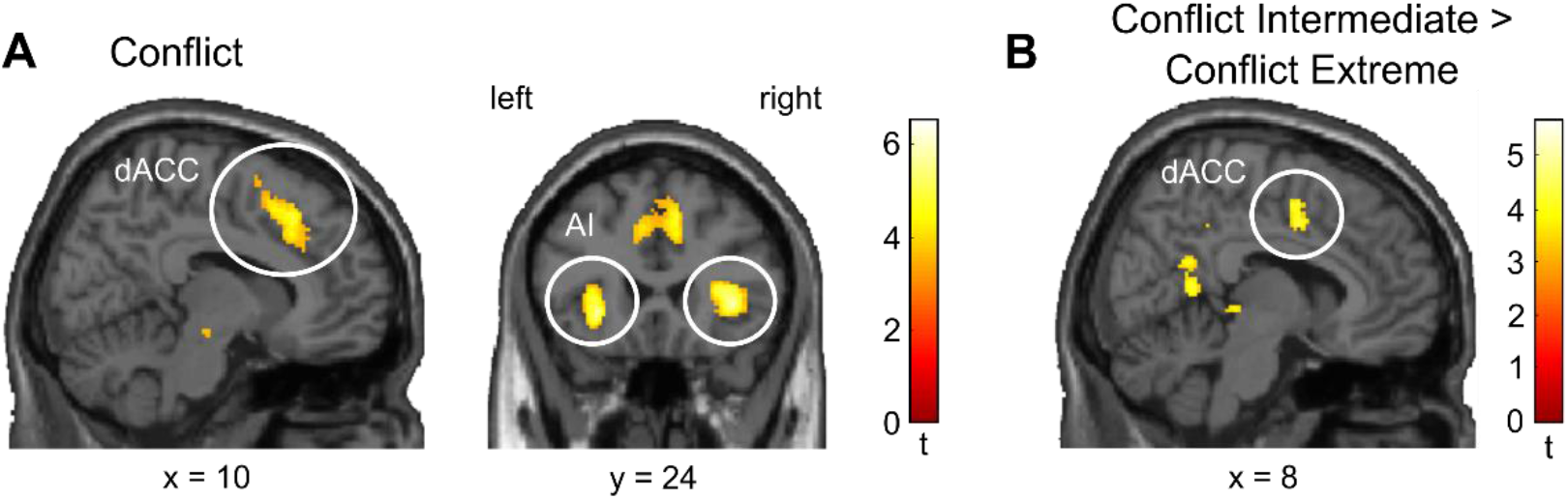
Effects of conflict. **(A)** BOLD-activity in dACC and bilateral anterior Insula (AI) correlated with conflict. **(B)** The correlation of BOLD-activity in dACC with conflict was significantly increased during intermediate versus extreme offers. Activations displayed at p < 0.001 uncorrected. See Table 1 for peak MNI-coordinates and statistics, significant at p<0.05 FWE corrected.

Next, we tested our main hypothesis that the extreme and intermediate conditions are treated as different control contexts by the brain. Note that one obvious reason for the effects in the categorical contrast Intermediate > Extreme could be that average conflict was higher in intermediate trials than in extreme trials (S3 Fig). Another possibility would be that the brain areas involved in processing conflict are modulated by the context. In other words, is conflict processed differently in the brain when the subject is in an intermediate compared to an extreme trial? Behavioural results in Fig. 3C already indicate such a context-dependent mechanism, showing that reaction times increased more with conflict during intermediate than extreme trials.

To test this context dependency using brain activity we included conflict as parametric modulator in our GLM, separately for intermediate and extreme offers. We then computed a contrast between the parametric modulator of conflict for intermediate offers minus the parametric modulator of conflict for extreme offers (Conflict Intermediate > Conflict Extreme). Our expectation was that the dACC would track conflicts (as a proxy for the value of planning), but to a lesser extent in a context with a low prior need for planning (i.e. in the extreme context), due to the metacosts associated with obtaining conflict values. We indeed found that BOLD-activity in dACC and right posterior middle temporal gyrus (pMTG) was more strongly correlated with conflict during intermediate offers compared to extreme offers (Fig 5B, Table 1 and S5 Table). An effect in dlPFC emerged at a lower threshold (see S5 Table). This finding aligns well with the results of the reaction time analysis and is consistent with the idea that the situation-appropriate investment into planning is driven by a context-dependent evaluation of conflict involving the dACC.

## Discussion

We used a novel sequential task with a complex task space to investigate how people decide when to plan ahead. We found evidence that participants use readily available features of the task space, such as offer values, to construct contexts that condition the balancing between forward planning and a simpler response strategy. We further provided evidence that the context-dependency of planning might be mediated by context-dependent conflict processing involving dACC. Our study provides initial evidence that the human ability to efficiently allocate cognitive control in complex tasks is supported by category-based cognition that harnesses regularities in control demand to generate control contexts.

Normatively, a decision about the engagement into elaborate planning should find the optimal tradeoff between the benefits and costs of such planning in a given situation (Shenhav et al., 2013). It is however unlikely that people calculate the expected value of planning explicitly because this would require planning itself. Previous studies have therefore argued that people must have quick and automatic access to a proxy of the value of planning (D. G. Lee et al., 2021; Lieder et al., 2018). It is a largely unresolved question how humans learn such proxies to decide about their engagement into effortful planning. A hallmark of learning is that people generalize individual sensorimotor experiences into broader categories allowing them to react adequately to novel instances of a learned category (Konidaris, 2019; E. E. Smith et al., 1981; Tenenbaum et al., 2011; Yee, 2019). Recent research suggests that the brain leverages similar generalisation mechanisms when deciding about the allocation of cognitive control or when computing meta-control (Bhandari et al., 2017; M. Botvinick et al., 2020; Lieder et al., 2018; Marković et al., 2020; Schwöbel et al., 2021). Here, we tested this principle using a complex planning task, where the demand for planning changed depending on the current situation as defined by a configuration of task features including offer value, energy, current energy cost, future energy cost and trial within a segment.

Our results suggest that participants used a generalized task representation, mapping clusters of states (contexts) to an approximated value of planning (e.g. conflict) to decide efficiently about the investment in planning. In particular, our results suggest that participants constructed two contexts, one for the extreme offers 1 and 4, associated with a low prior tendency for planning, and one for the intermediate offers 2 and 3 associated with a high prior tendency for planning. Three pieces of converging evidence supported this conclusion: First, responses in an intermediate context were more mixed between accepting and rejecting compared to an extreme context. Second, our modelbased analysis suggested that the mixed response profile for the intermediate offers can be explained by participants planning ahead multiple steps into the future (especially offer 3, see Fig 2C). Third, the findings that the correlation of conflict with response times and dACC activity depended on the current context further corroborated that participants had learned an offer-specific context structure.

To investigate the usage of forward planning or simple heuristics and its determining factors, we designed a task where both types of decision making can occur. Our computational analysis indeed revealed that a model including both, planning and simple offer-specific preferences, fitted behaviour substantially better, than a model that relied on planning or a simple strategy alone. Our model implemented planning as a search through a decision tree, calculating expected long-term state-action values. However, due to the high computational cost of such an exhaustive search, it is unlikely that participants planned in exactly this way. We therefore assumed that participants used some sort of approximate planning, reducing planning complexity while still accommodating for an action’s future consequences. Pruning of decision trees (Huys et al., 2012), adjusting the planning depth (Keramati et al., 2016) or the division of a decision problem into smaller subproblems (Huys et al., 2015) are just some ways of how people can adjust planning complexity. While we cannot say anything about the exact implementation of planning, we found that the fitted individual planning weights of the winning hybrid model correlated with RT, suggesting that participants engaged in some kind of forward planning. Another limitation of the hybrid model is that it does not explicitly model arbitration between different decision modes, but only recognises the presence of a mixture. While research on model-free and model-based reinforcement learning systems provided important insights into the neurocomputational mechanism of arbitration using a simple (two-step) planning task (Kool et al., 2017; S. W. Lee et al., 2014), less is known about how the brain adapts its decision mode during more complex tasks. Our study provides evidence for a link between the generalisation of task representations and high-level control decisions and may therefore inform future attempts to model arbitration mechanisms in more complex realistic environments.

Our finding that BOLD activity in the dACC correlated with conflict is consistent with the view that the dACC monitors the need for effortful controlled processing (M. M. Botvinick et al., 2001; Shenhav et al., 2013; Shenhav et al., 2017). In our task, when it was not clear at the beginning of a trial whether accepting or rejecting is the best option (i.e. if the conflict was high), participants needed to generate additional information by planning ahead which could then help to adjudicate between the two options. While dACC might have played a central role in detecting the need for additional planning, a distributed network including dlPFC and other structures might have been involved in the additional information sampling. The finding that dlPFC was more active in the demanding intermediate context is consistent with the view that dlPFC is central to planning (Fuster, 2015). In order for planning processes to have an impact on the decision, it is often necessary to inhibit a prepotent response first. We found evidence that such prepotent responses played a role in in our task as well, as our model-based analysis revealed non-negative choice preference parameters for all offers. Previous research suggests that such prepotent responses could have been inhibited by a hyper-direct pathway from the ACC to the basal ganglia, effectively increasing time until movement generation and thus allowing planning structures to influence decision making (Cavanagh et al., 2011; Gluth et al., 2012; Wiecki & Frank, 2013; Wiecki, Sofer, et al., 2013).

The anterior insula (AI) is often coactivated with the ACC in classical cognitive control tasks (Duncan et al., 2000) and sequential tasks alike (e.g. Schwartenbeck et al., 2015). Previous research suggests that, besides of its role in interoception and general awareness (Craig et al., 2009), the AI appears to be specifically involved in the representation and learning of uncertainty (Bossaerts, 2010; Loued-Khenissi et al., 2020; Singer et al., 2009). Translated to our task, the AI could have relayed information about uncertainty (or conflict) to the dACC, which then initiates adaptive behavioural change in the form of planning.

We found that the correlation of activity in the dACC with conflict (which we take to be a proxy for the value to plan ahead) depended on context. One possible explanation for this pattern could be that the dACC has access to a hierarchical representation of learned conflicts, whereby conflicts encoded at a finer level of task space are subsumed under conflicts encoded at the level of context. In other words, states of similar difficulty could be grouped into a more general category that e.g. simply indicates whether the decision is easy or difficult. In contexts with a high prior expectation of conflict, i.e. in an intermediate context, the dACC could access conflict at a more fine-grained level to enable the appropriate level of planning. Conversely, in a context with low prior expectation of conflict, i.e. in an extreme context, the dACC would not access information beyond that at the coarse context level, as the overall need for planning was low anyway. Speculating on the algorithmic implementation of such a process, the context-dependent prior assumption about conflict could set the threshold for the meta-decision problem of inferring the need of planning. In an intermediate context, a high meta-threshold would grant enough time for a state-level readout of conflict, whereas in an extreme context the need for planning would have been determined before state-level conflicts were accessed. We also found evidence that right posterior middle temporal gyrus (pMTG) is more correlated with conflict in an intermediate than in an extreme context. Previous research implicated the pMTG in category-based cognition (Martin, 2007). It is therefore an intriguing possibility that the pMTG is also capable of forming abstract categories of choice difficulty that support the context-dependent evaluation of planning demands. Although we can only speculate about the role of pMTG, the question how brain mechanism for structured knowledge acquisition and cognitive control interact is an important direction for future research. Overall, our findings are generally consistent with the view that people exploit the structure of a task for efficient storage and access of the value of control (Lieder et al., 2018).

## Acknowledgements

The authors thank Florian Bolenz and Philipp T. Neukam for valuable comments and suggestions. This work was funded by the German Research Foundation (DFG, Deutsche Forschungsgemeinschaft), SFB 940 - Project number 178833530, TRR 265 - Project number 402170461 and was partially supported by Germany’s Excellence Strategy – EXC 2050/1 – Project number 390696704 – Cluster of Excellence “Centre for Tactile Internet with Human-in-the-Loop” (CeTI) of Technische Universität Dresden.

## Supplementary information

**S1 Table.**
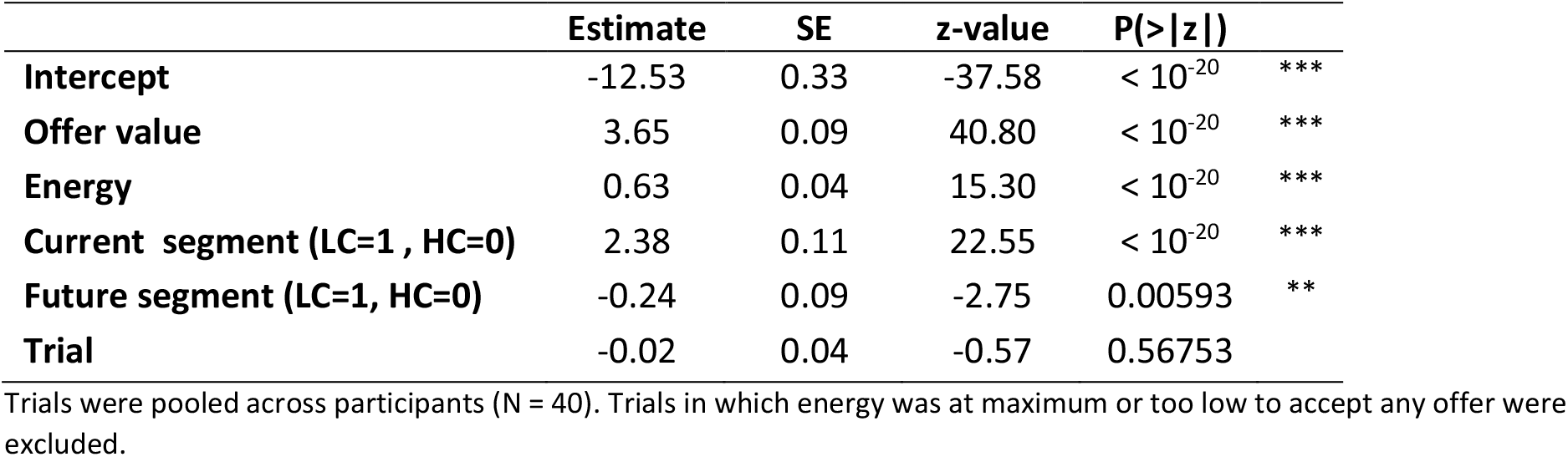
Logistic regression of choice (accept = 1, reject = 0) against task features.

|  | Estimate | SE | z-value | P(>\|z\|) |  |
| --- | --- | --- | --- | --- | --- |
| **Intercept** | -12.53 | 0.33 | -37.58 | < 10<sup>-20</sup> | *** |
| **Offer value** | 3.65 | 0.09 | 40.80 | < 10<sup>-20</sup> | *** |
| **Energy** | 0.63 | 0.04 | 15.30 | < 10<sup>-20</sup> | *** |
| **Current segment (LC=1, HC=0)** | 2.38 | 0.11 | 22.55 | < 10<sup>-20</sup> | *** |
| **Future segment (LC=1, HC=0)** | -0.24 | 0.09 | -2.75 | 0.00593 | ** |
| **Trial** | -0.02 | 0.04 | -0.57 | 0.56753 |  |

Trials were pooled across participants (N = 40). Trials in which energy was at maximum or too low to accept any offer were excluded.

**S2 Table.** fMRI results for the contrast: intermediate > extreme.

| Region | cluster size<br>(voxels) | t | FWE p value |  | MNI coordinates |  |  |
| --- | --- | --- | --- | --- | --- | --- | --- |
|  |  |  | whole brain | SVC | x | y | z |
| Dorsal anterior cingulate | 929 | 7.26 | <0.001 | <0.001 | -4 | 16 | 52 |
| Occipital L | 1252 | 6.98 | <0.001 |  | -18 | -92 | -10 |
| Occipital R | 1475 | 6.38 | 0.007 |  | 20 | -84 | -8 |
| Dorsolateral prefrontal R | 106 | 5.12 | 0.180 | 0.018 | 52 | 34 | 24 |
| Superior parietal L | 76 | 4.68 | 0.456 |  | -28 | -64 | 52 |
| Putamen R | 95 | 4.64 | 0.486 |  | 20 | 16 | -4 |
| Angular R | 112 | 4.54 | 0.580 |  | 34 | -70 | 44 |
| Occipital L | 92 | 4.51 | 0.602 |  | -24 | -84 | 14 |
| Anterior Insula L | 45 | 4.09 | 0.918 |  | -26 | 18 | 0 |
| Dorsolateral prefrontal L | 30 | 4.06 | 0.935 | 0.223 | -48 | 32 | 24 |
| Precentral L | 21 | 3.98 | 0.961 |  | -36 | -12 | 50 |
| Inferior parietal R | 50 | 3.97 | 0.964 |  | 36 | -48 | 38 |
| Superior frontal L | 11 | 3.71 | 0.997 |  | -24 | 40 | -14 |
| Precuneus R | 128 | 3.69 | 0.998 |  | 6 | -62 | 34 |

The table contains all clusters with more than 10 voxels that survived uncorrected statistical thresholding with p < 0.001.

**S3 Table.** fMRI results for the contrast: extreme > intermediate.

| Region | cluster size<br>(voxels) | t | FWE p value |  | MNI coordinates |  |  |
| --- | --- | --- | --- | --- | --- | --- | --- |
|  |  |  | whole brain | SVC | x | y | z |
| Posterior parietal L | 483 | 6.68 | 0.003 |  | -60 | -44 | 38 |
| Posterior parietal R | 439 | 5.49 | 0.072 |  | 64 | -32 | 28 |
| Posterior cingulate | 106 | 5.15 | 0.169 |  | 6 | -6 | 36 |
| Amygdala L | 186 | 4.92 | 0.280 |  | -24 | -2 | -18 |
| Middle temporal L | 391 | 4.90 | 0.296 |  | -48 | -64 | 6 |
| Middle temporal R | 237 | 4.86 | 0.323 |  | 48 | -12 | -12 |
| Amygdala R | 94 | 4.56 | 0.560 |  | 28 | -2 | -18 |
| Ventral striatum | 23 | 4.40 | 0.701 |  | -4 | 22 | -10 |
| Middle temporal R | 109 | 4.20 | 0.858 |  | 66 | -48 | 10 |
| Anterior cingulate | 16 | 4.09 | 0.920 |  | -4 | 32 | 4 |
| Anterior Insula R | 20 | 3.87 | 0.984 |  | 30 | 20 | -16 |
| Superior frontal R | 17 | 3.71 | 0.997 |  | 10 | 48 | 26 |

The table contains all clusters with more than 10 voxels that survived uncorrected statistical thresholding with $p < 0.001$ .

**S4 Table.** fMRI results for parametric effect of conflict.

| Region | cluster size<br>(voxels) | t | FWE p value |  | MNI coordinates |  |  |
| --- | --- | --- | --- | --- | --- | --- | --- |
|  |  |  | whole brain | SVC | x | y | z |
| Anterior Insula R | 523 | 6.50 | 0.004 |  | 38 | 20 | 0 |
| Anterior Insula L | 600 | 5.62 | 0.041 |  | -30 | 24 | -2 |
| Dorsal anterior cingulate | 644 | 4.82 | 0.290 | 0.007 | 10 | 24 | 44 |
| Thalamus R | 40 | 4.29 | 0.718 |  | 8 | -18 | -12 |
| Pallidum L | 29 | 4.01 | 0.909 |  | -12 | 0 | -4 |
| Supplementary Motor Area R | 24 | 3.78 | 0.983 |  | 12 | 8 | 62 |

The table contains all clusters with more than 10 voxels that survived uncorrected statistical thresholding with $p < 0.001$ .

**S5 Table.** fMRI results for the contrast: conflict intermediate > conflict extreme.

| Region | cluster size<br>(voxels) | t | FWE p value |  | MNI coordinates |  |  |
| --- | --- | --- | --- | --- | --- | --- | --- |
|  |  |  | whole brain | SVC | x | y | z |
| Middle temporal R | 489 | 5.69 | 0.040 |  | 56 | -64 | 6 |
| Anterior Insula R | 389 | 5.05 | 0.196 |  | 36 | -8 | 18 |
| posterior cingulate L | 29 | 4.79 | 0.345 |  | -16 | -32 | 40 |
| Occipital L | 441 | 4.75 | 0.378 |  | -48 | -74 | 24 |
| Precuneus R | 117 | 4.54 | 0.551 |  | 10 | -50 | 10 |
| Dorsolateral prefrontal R | 208 | 4.49 | 0.592 | 0.083 | 54 | 28 | 6 |
| Angular R | 50 | 4.41 | 0.667 |  | 38 | -62 | 26 |
| Precuneus R | 50 | 4.36 | 0.712 |  | 12 | -44 | 40 |
| Dorsal anterior cingulate | 105 | 4.32 | 0.744 | 0.03 | 8 | 2 | 40 |
| Cuneus L | 91 | 4.28 | 0.774 |  | -10 | -68 | 26 |
| Superior temporal R | 32 | 4.16 | 0.859 |  | 46 | -42 | 22 |
| Thalamus L | 32 | 4.08 | 0.905 |  | -6 | -30 | -2 |
| Occipital L | 35 | 4.03 | 0.929 |  | -24 | -86 | 38 |
| Occipital L | 26 | 3.93 | 0.966 |  | -16 | -48 | 0 |
| Superior temporal R | 19 | 3.90 | 0.971 |  | 44 | -14 | -10 |
| Precentral R | 117 | 3.83 | 0.984 |  | 40 | -18 | 58 |
| Superior temporal L | 44 | 3.72 | 0.995 |  | -54 | -20 | 12 |
| Thalamus R | 20 | 3.72 | 0.995 |  | 8 | -30 | -2 |
| Precuneus R | 60 | 3.71 | 0.996 |  | 16 | -60 | 26 |
| Supplementary motor area R | 21 | 3.69 | 0.997 |  | 4 | -8 | 58 |
| Middle temporal L | 12 | 3.69 | 0.997 |  | -62 | -52 | 0 |

The table contains all clusters with more than 10 voxels that survived uncorrected statistical thresholding with $p < 0.001$ .

**S1 Figure.**
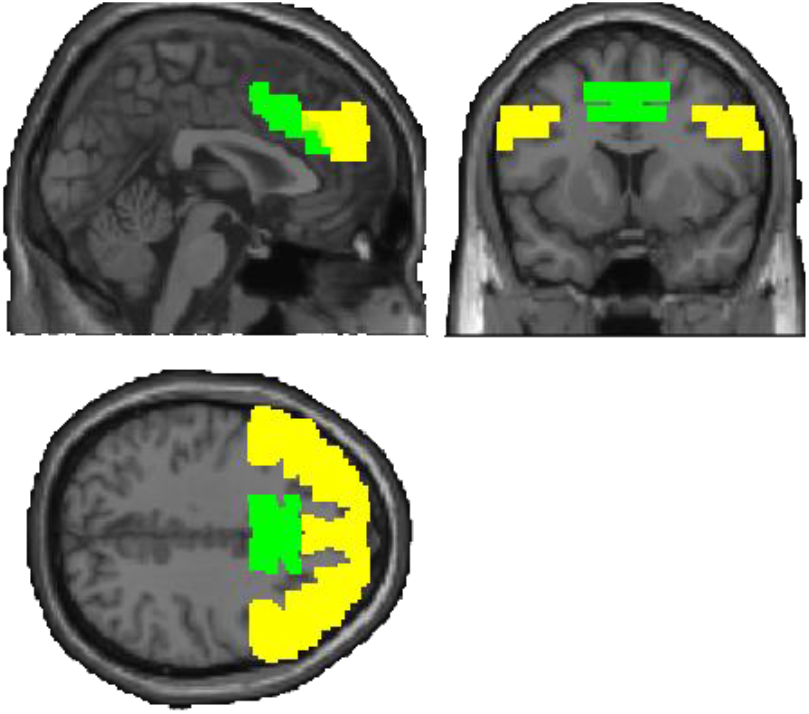
Definition of anatomical ROIs for small volume correction. dACC (green), dlPFC (yellow).

**S2 Figure.**
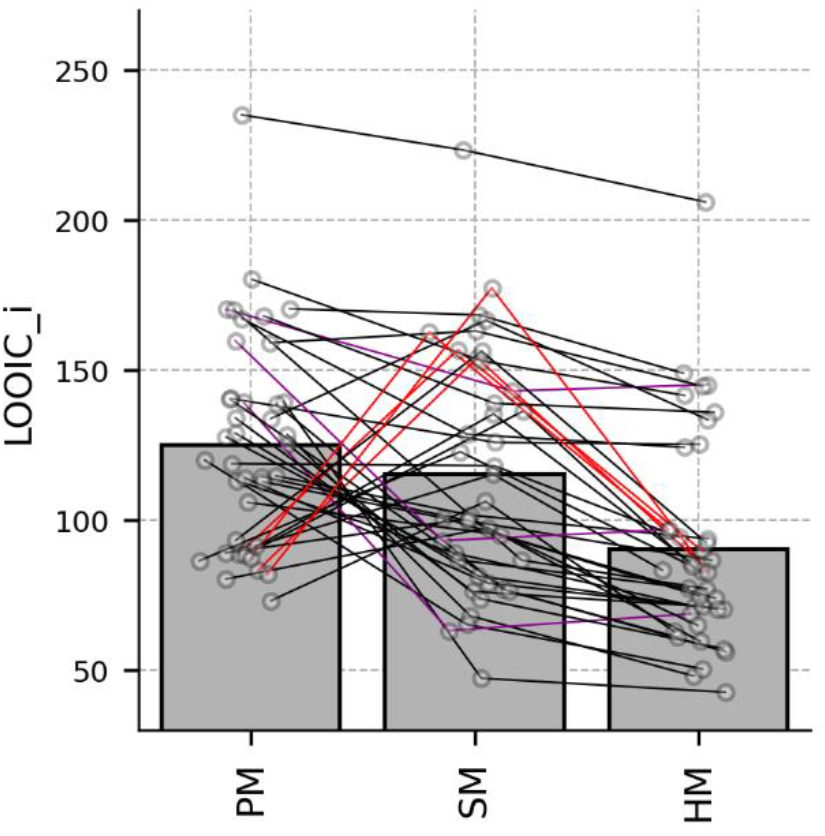
Comparing models on the participant level. To calculate the predictive accuracy for each participant (LOOIC_i), pointwise log predictive densities (see methods) have to be summed up for all trials within each participant (white dots). Note that this differs from Fig 2B, in which the individual pointwise log predictive densities are summed up over all trials and participants. The hybrid strategy model (HM) was best for 33 (of N = 40) participants (black lines). For 3 participants the simple strategy model (SM, violet lines) and for 4 participants the planning model (PM, red lines) explained behaviour best. Bars indicate average LOOIC_i.

**S3 Figure.**
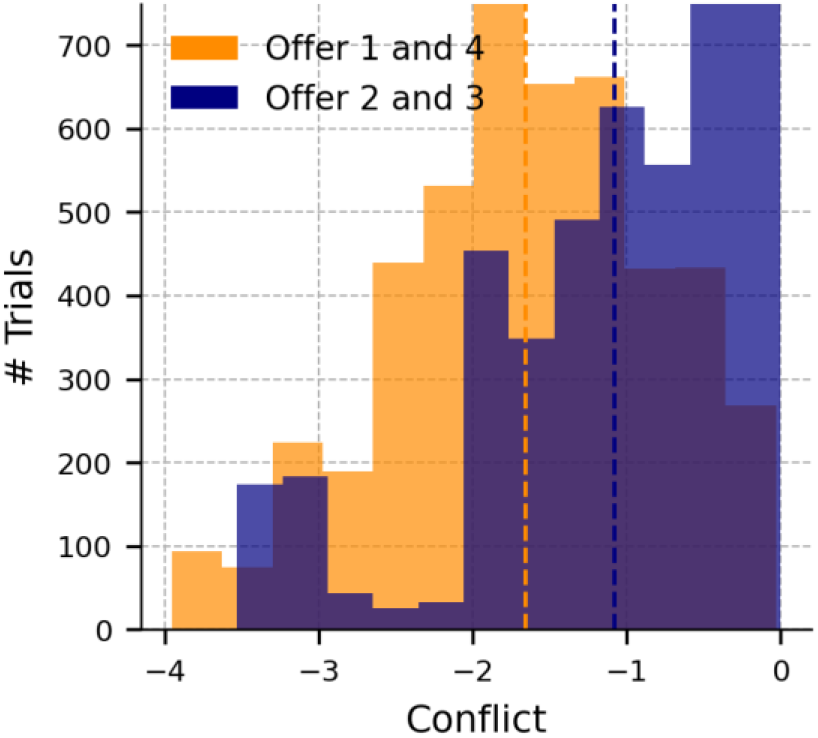
Encountered conflict levels plotted for intermediate (2, 3) and extreme (1, 4) offers. Histograms contain data of all forty participants. Dashed lines indicate the mean.

### S1 Text. Task instructions

Participants were guided step by step through the task on the computer screen. Written instructions were accompanied by an arrow pointing towards the stimulus element currently addressed. The written instructions were:

♦ Dear participant, the experiment is about collecting as many points as possible. Depending on your score at the end of the experiment, you will be paid a monetary bonus.
♦ The current score is indicated by the upper yellow bar.
♦ You can get points by accepting offers.
♦ The current offer is presented in the middle of the screen. The magnitude of offers varies between 1 and 4, represented by the number of golden trophies. The offers are drawn at random, having the same occurrence probability of 25%.
♦ However, accepting an offer is associated with energy costs. Your current energy level is represented by the lower blue bar. If you accept an offer and do not have enough energy, no points will be credited to you and the next trial will begin.
♦ You can replenish your energy account by selecting the “reject” option. This will increase your energy level by 1 and the next trial will begin.
♦ The energy level can have a maximum value of 6.
♦ The experiment is divided into segments, each consisting of 4 trials. Two numbers are displayed on the screen to indicate how far you are in the current segment.
♦ There are 2 different segment types, in which the energy costs for accepting an offer differ. In segments with 1 flash, 1 energy unit is subtracted when you accept an offer. In segments with 2 flashes, 2 energy units are subtracted when you accept an offer. The left blue-orange box at the bottom right of the screen informs you about the type of the current segment.
♦ In addition to the type of the current segment, information about the energy costs in the next segment is available. This can be seen in the right blue-orange box at the bottom right of the screen.
♦ Breaks: During the main experiment in the scanner, you have the possibility to pause twice. The pause screen is automatically displayed. You decide when you are ready to continue the experiment. Note: After a pause your score will be reset to 0. This has no effect on your final bonus. Your score is counted continuously.
♦ Deadline: You have a maximum of 5 seconds for each decision. If you exceed this time limit, the next trial will begin without points being awarded.
♦ Training: Before the main experiment in the scanner starts, you will be given a few training trials on the PC to familiarize yourself with the experiment. There is no deadline in the training and the points gained here have no effect on the bonus paid out. Please try to get as many points as possible anyway. The training phase will end automatically.

